# Automated AI image recognition tools improve the efficiency of aerial wildlife counts: A multi-species case study on breeding seabirds and pinnipeds at the sub-Antarctic Bounty Islands

**DOI:** 10.64898/2026.02.14.705878

**Authors:** Chris G. Muller, Ryan King, G. Barry Baker, Katrina Jensz, Fred Samandari

## Abstract

Accurate monitoring of populations is essential for conservation management, including for vulnerable seabirds. Yet traditional ground-based surveys are logistically challenging and time-consuming, especially in remote environments such as the sub-Antarctic islands. Advances in aerial imagery and artificial intelligence (AI) offer opportunities to improve the efficiency and repeatability of population surveys. In this study, we evaluate an AI-based approach for counting Salvin’s albatross from high-resolution aerial imagery collected using a piloted fixed-wing aircraft at the Bounty Islands, New Zealand. Imagery acquired during a single-day survey was processed to create orthomosaic images, which were previously analysed using manual counts by an experienced observer. We applied an automated detection and counting model based on a Faster R-CNN architecture with Slicing-Aided Hyper-Inference, and compared AI-derived counts with original human counts in terms of accuracy, consistency, and processing time. The AI achieved an initial F1 score of 92.8% for albatross detection and produced counts within 3% of the manual results, while reducing processing time from approximately 66 hours to just over four minutes. The model was also capable of simultaneously detecting additional species present within the mixed breeding colony, including erect-crested penguins, fulmar prions, and New Zealand fur seals, adding scalable efficiency gains for future surveys. Our results demonstrate that combining piloted aircraft surveys with AI-based image analysis provides a rapid, scalable, and accurate method for monitoring seabird populations, with substantial benefits for conservation management in remote and logistically constrained regions.

## Introduction

Accurate estimation of seabird population size is fundamental to conservation management, particularly for long-lived, wide-ranging species such as albatrosses which are exposed to multiple natural and anthropogenic threats across their life cycle (Dias et al., 2019; Paleczny et al., 2015; Tuck et al., 2015). Traditional ground-based counts of breeding colonies are often logistically challenging, costly, and potentially disruptive, especially in remote sub-Antarctic environments where access is constrained by weather, terrain, and biosecurity considerations (Baker et al., 2020; Baker & Jensz, 2019; Parker et al., 2020; Parker et al., 2016).

Aerial monitoring of wildlife is increasingly being used to take advantage of data sourced from satellites, piloted aircraft, and uncrewed aerial vehicles (UAVs), with data collection including standard cameras, thermal imagery, and automated mapping of VHF transmitters (Corcoran et al., 2021; Lynch et al., 2012; Muller et al., 2019). As a result, aerial population survey techniques have become an increasingly important component of seabird monitoring programmes, enabling comprehensive coverage of colonies while reducing time spent in the field (Arata et al., 2003; Fretwell et al., 2017). However, manual counting of animals from collected imagery can be time-consuming, and prone to errors, particularly with inexperienced counters (Chrétien et al., 2016; Jones et al., 2023; Meek et al., 2012).

Recent advances combining high-resolution aerial imagery with machine learning have transformed wildlife survey methods by enabling automated detection and counting of animals from imagery that would otherwise require extensive manual processing (Corcoran et al., 2021). Deep learning approaches, in particular convolutional neural networks, have demonstrated strong performance in detecting and enumerating birds and other wildlife across a range of platforms (Borowicz et al., 2019; Gaurav et al., 2024). For seabirds, these approaches have been successfully applied to species ranging from penguins to albatrosses, offering improvements in efficiency, repeatability, and transparency relative to manual image interpretation (Borowicz et al., 2019; Bowler et al., 2020; Fretwell et al., 2017).

The Bounty Islands (Fig. 1) lie approximately 700 km (east-south-east of the South Island of New Zealand (LINZ, 2012), requiring several days’ travel to reach by sea.

**Fig 1.**
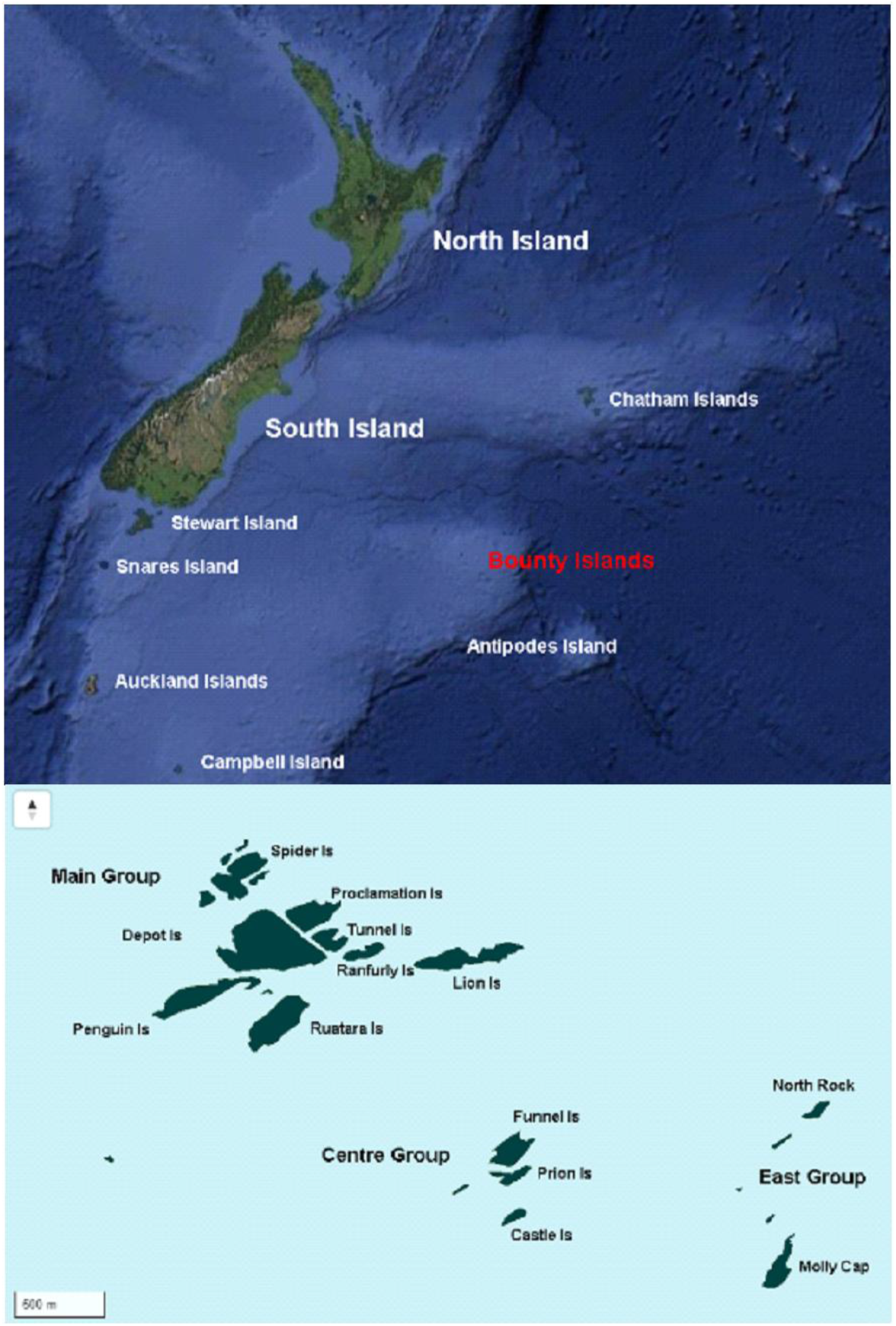
Map New Zealand and sub-Antarctic islands (upper panel) showing the location of the Bounty Islands in red (Basemap from Google Maps). Detail (lower panel) showing islands in the Bounty Islands archipelago (Basemap from LINZ).

The archipelago consists of 13 separate granite islands, with a total land area of 1.35 km^2^ (Department of Conservation, 2026). The rocky islands have few safe landing points, and there is virtually no soil or vegetation with the rocks subjected to frequent large waves and salt spray, and much of the island is occupied by breeding wildlife making access difficult for ground research (CGM pers. obs). Within New Zealand, the Bounty Islands represent one of the most important breeding sites for Salvin’s albatross (*Thalassarche salvini*), a species classified as Vulnerable due to ongoing bycatch in commercial fisheries and its restricted breeding distribution (IUCN, 2025). The Bounty Islands are also the primary breeding area for erect-crested penguins (*Eudyptes sclateri*), fulmar prions (*Pachyptila crassirostris*), and one of the largest breeding colonies of New Zealand fur seals (*Arctocephalus forsteri*), estimated at around 20,000 individuals (Department of Conservation, 2026). Together with Salvin’s albatross, these species share a mixed multi-species breeding area (Baker & Jensz, 2019). However, surveys of penguin populations on the Bounty Islands have previously relied on manual ground-based counts (Hiscock & Chilvers, 2014).

Previous albatross population surveys have demonstrated the feasibility of using aerial imagery with manual counts from high-resolution photographs to provide population estimates for Salvin’s albatross (Baker & Jensz, 2019), and also other seabird species in the New Zealand sub-Antarctic (Baker et al., 2020; Parker et al., 2017), while at other locations satellite imagery has also been used to survey albatross species (Arata et al., 2003). More recently, deep learning approaches have been developed to automate the detection and counting of Salvin’s albatrosses from UAV-derived imagery, showing that AI-based methods can be a useful tool when appropriately developed, trained and validated (Bowler et al., 2020; Rogers et al., 2025).

However, most AI-based seabird surveys to date have relied on imagery collected either from satellites, which can have issues with poor resolution making target recognition difficult (Bowler et al., 2020); or from UAV platforms which can also be limited in spatial resolution as a result of using smaller, lighter cameras which they are able to carry (Rogers et al., 2025). Smaller UAVs can also be limited in their flight endurance and therefore survey area coverage, reducing operational flexibility and their efficiency when operating in isolated field sites. In contrast, piloted fixed-wing aircraft provide an alternative platform capable of rapidly surveying large and remote colonies in a single flight, while capturing imagery of improved resolution allowing clearer identification of individual birds (Baker & Jensz, 2019). To date, the application of modern AI-based counting methods to imagery collected from piloted aircraft remains comparatively under-explored, particularly in the New Zealand sub-Antarctic, despite the extensive historical archives of such data and their continued use in wildlife monitoring.

In this study, we evaluate the use of an automated AI-based recognition and counting tool to survey Salvin’s albatrosses from high-resolution aerial imagery collected from a piloted aircraft. Building on established aerial survey and image-processing workflows, we compare automated counts directly with previously completed manual image counts undertaken by an experienced observer, and we quantify differences in accuracy, consistency, and processing time. A secondary aim was to improve conservation management opportunities by exploring simultaneous surveys of multiple wildlife species at the same time.

By integrating AI-based analysis with proven aerial survey methods, this work aims to demonstrate a scalable and efficient approach to seabird and pinniped population monitoring with tools supporting piloted aircraft surveys, and which are also compatible with existing UAV- and satellite-based techniques. This flexible approach enables timely and accurate conservation decision-making for threatened species.

## Methods

### Fieldwork

Fieldwork was carried out at the Bounty Islands (47°45′S 179°03′E) in the New Zealand sub-Antarctic (Fig. 1). Surveys of nesting Salvin’s albatrosses were conducted on 25 October 2018, using a twin turboprop Reims F406 aircraft. The aircraft departed from Dunedin airport on mainland New Zealand, then returned to base on the same day. Survey timing was aimed to coincide with the middle of the day to maximise available daylight for good visibility, and minimise the number of loafing non-breeders likely to be present (Baker et al., 2018). The flight was scheduled during mid-October to overlap with the peak incubation period of the albatross breeding cycle, as well as the presence of a scientific field team on the islands who were able to provide ground-truthing data for the original survey counts (Baker & Jensz, 2019).

Surveys were conducted at an altitude of 1200 ft AGL, and a flight speed of 120 kt, consisting of a series of parallel transects across the island group. Transects had a 100 m separation distance with 20% side overlap and 45% frontal overlap on imagery For details see (Baker & Jensz, 2019). Images were taken using a full-frame Digital Single Lens Reflex Camera (Nikon D800) and telephoto lens (Nikon 70-200 mm f2.8) set at 70mm photo extension. Shutter speed was set to 1/2500 sec, and operated in shutter priority mode to maintain shutter speed. The camera was mounted through a porthole in the fuselage to collect nadir imagery (zero degree deviation from vertical) without affecting aircraft streamlining. Use of a dedicated camera operator freed up the pilot for flying and navigation tasks. Photographs were saved as high-resolution JPEG format files for later analysis.

### Manual Counting

Individual images were stitched together using Adobe Photoshop software (www.Adobe.com) to create ortho-mosaic images, using methods established for previous aerial albatross surveys (Arata et al., 2003; Baker et al., 2018). Photographs were stitched in sequence to construct photo-montages of each transect flown. From these, a complete series of overlapping images were generated that covered the entire albatrosses nesting area for each island in the archipelago.

To perform manual counting, images were viewed in Photoshop and magnified to allow identification of birds. All Salvin’s albatrosses on each montage were identified and marked with a coloured circle using the Paintbrush tool to prevent double-counting. To assist with counting, Mousecount software (mousecount.software.informer.com) and a hand held click counter were also used. For details see (Baker & Jensz, 2019).

### AI Counting

We repeated the recognition and counting phase using an automated AI software tool utilising machine learning tools. These automated counts were then compared with the original results counted by a human observer. The AI detection algorithm was created to automatically recognise and count animals from image files, and utilised a Faster-RCNN model with a Residual Network backbone and a Feature Pyramid Network. Inference was conducted using the Slicing-Aided Hyper-Inference (SAHI) method to improve detection for images with large number of individuals. The model was trained using a nominal 1000 examples of the primary target species, and opportunistically trained on secondary species (including erect-crested penguins, fulmar prions (referred to as petrels), and fur seals) using whatever additional data were available in the primary training images. Secondary species therefore received less than the expected amount of training because there were fewer individuals present in the training images.

Accuracy of the model was measured using F1 scores to collate false positive and false negative detections, thereby measuring the precision and recall scores. F1 scores provide a more accurate measure of accuracy than comparing output, as detection counts could otherwise be masked by detection inaccuracies (Jones et al., 2023; Meek et al., 2012). Verification testing was carried out using a similar-sized dataset to the initial training data, with approximately 1000 examples of the primary target species (albatross). F1 scores and a confusion matrix were created using a pre-existing set of annotations for the validation images. These were algorithmically compared to the detections predicted by the model for the same images, with an Intersection over Union (IoU) score of >0.5 between a predicted bounding box and validation box of the same category counting as a true positive. The validation dataset was split by selecting 65% of image tiles for training and 45% for validation. This proportion was chosen in order to test the model output with a smaller amount of available training data.

## Results

The aerial survey of the Bounty Islands was conducted between 1434–1545 hrs, taking 71 minutes. Total time in the air was from 1245–1745 hrs, a flight time of 5 hours. A total of 450 individual photographs were stitched together to create nine ortho-mosaic images; representing each island in the archipelago. The human observer counted 30,899 albatross, annotating each with a yellow dot (Fig. 2).

**Fig 2.**
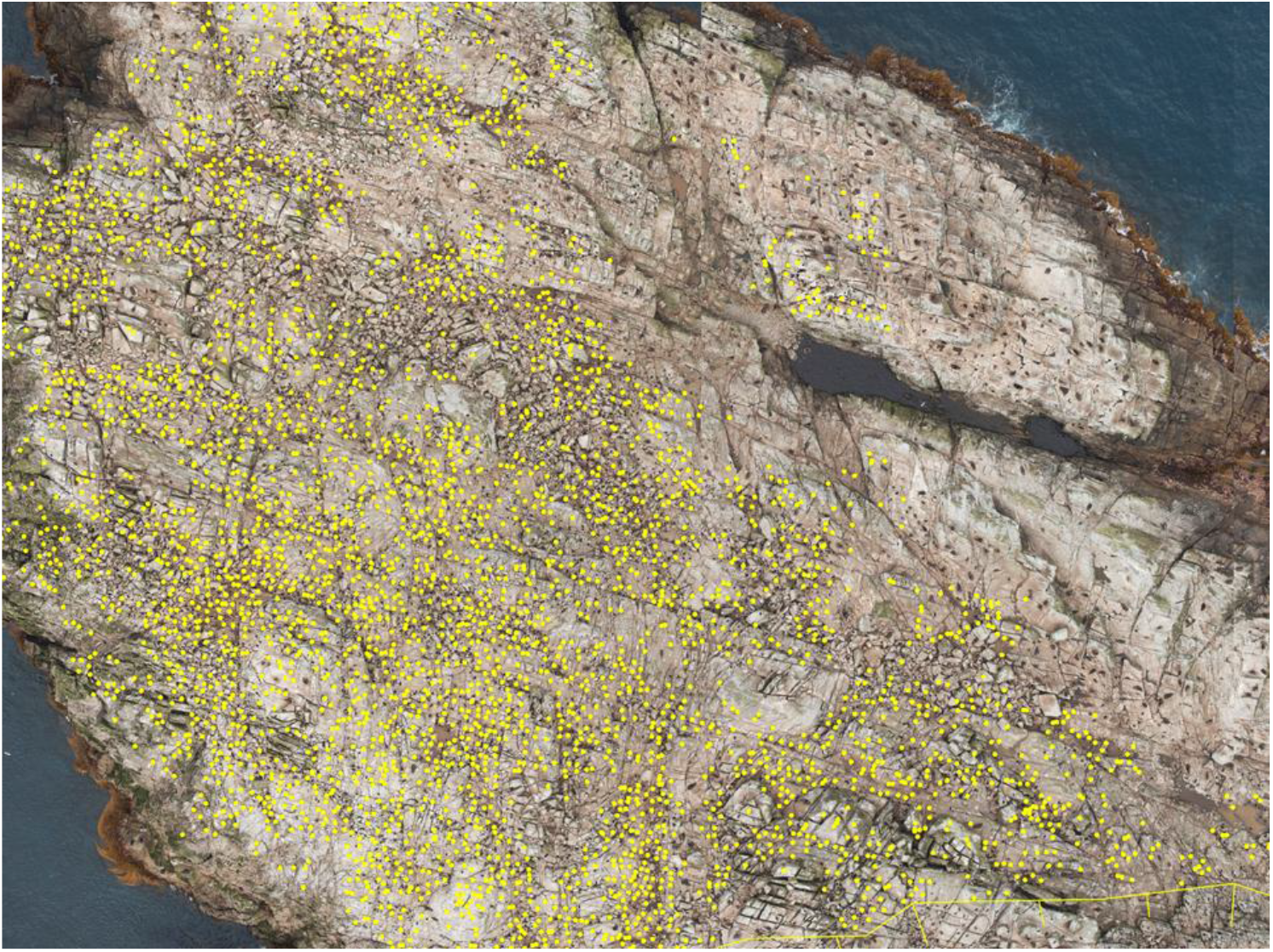
Cropped image showing examples of human counts; Salvin’s albatrosses are marked with a yellow dot.

After training, the AI model provided an F1 score representing an initial detection accuracy of 92.8% for the target species (Salvin’s albatross), with decreasing initial accuracy figures for each secondary species (Table 1).

**Table 1.**
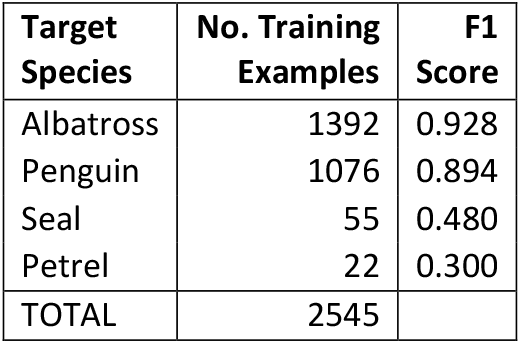
Training data summary showing the number of examples of each species used for training and verification of the model. F1 scores show the initial accuracy of AI output for the primary target (albatross), and secondary target species. Only the primary target species included the expected amount of training data.

Following initial training, the verification testing showed that from a total of 1410 animals present in the test images, nine albatross were misidentified as other species, and 16 other species were misidentified as albatross (Table 2).

**Table 2.** Confusion matrix output from AI verification testing. The trained model was run on a small subset of images (n=1410), and output was confirmed by a human observer. The matrix shows correct identifications (green highlights), and incorrect detections comprising false positive and false negative detections (grey highlights).

| Annotation Category | Detected Category |  |  |  | Error |
| --- | --- | --- | --- | --- | --- |
|  | albatross | penguin | seal | petrel |  |
| albatross | 855 | 1 | 8 | 0 | 9 |
| penguin | 14 | 484 | 18 | 0 | 32 |
| seal | 0 | 0 | 23 | 0 | 0 |
| petrel | 2 | 2 | 0 | 3 | 4 |
| **Error** | 16 | 3 | 26 | 0 | **45** |

Following successful verification, around half of the original images were then presented to the AI for re-counting, which took 246 seconds (4.1 minutes) to count 31,718 albatrosses (Table 3, Fig. 3). In comparison, the human observer counted 30,899 albatrosses in the re-counted images (Table 3), a difference of 819 birds. The original human counts took 66 hours, or around 1.5 standard working weeks, making the AI counts 963 times faster.

**Table 3.** Count comparison of Salvin’s albatross at the Bounty Islands showing the difference between human counts and the AI re-count, including an accuracy measure.

| Location | Researcher Count | AI Re-count | Difference | Accuracy |
| --- | --- | --- | --- | --- |
| Molly Cap 1 | 3,728 | 3,651 | -77 | 98% |
| Molly Cap 2 | 1,474 | 1,393 | -81 | 95% |
| Funnel Is 2 | 4,633 | 4,455 | -178 | 96% |
| Funnel Is 3 | 85 | 82 | -3 | 96% |
| Funnel Is 5 | 1,137 | 1,115 | -22 | 98% |
| Main Group 3b | 2,991 | 4,104 | 1,113 | 63% |
| Main Group 4b | 11,757 | 11,803 | 46 | ~100% |
| Main Group 2 | 789 | 793 | 4 | 99% |
| Main Group 3a | 4,305 | 4,322 | 17 | 100% |
| **TOTAL** | **30,899** | **31,718** | **819** | **97%** |

**Fig 3.**
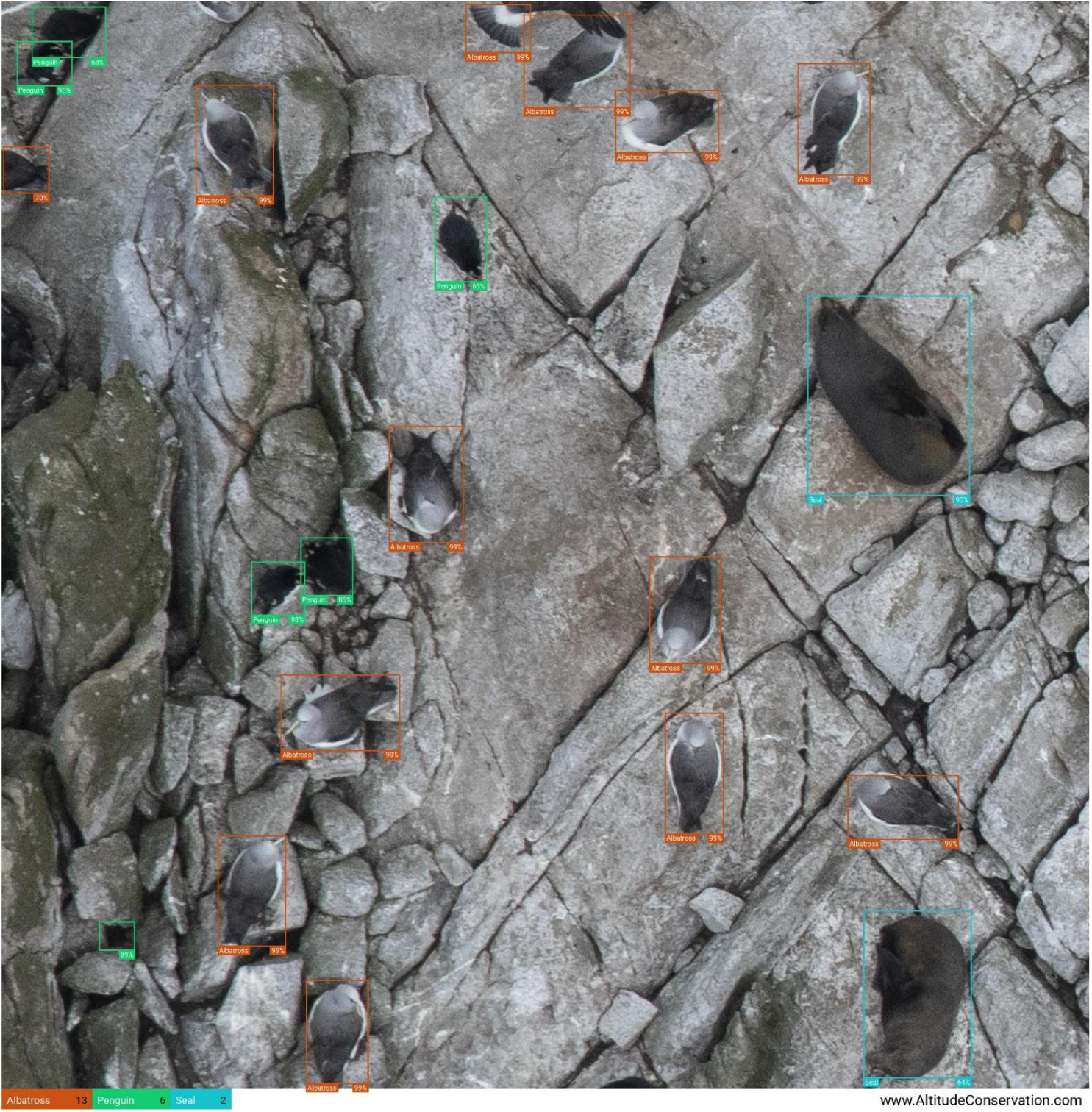
Zoomed image of AI output showing identified instances of all trained species, annotated with bounding boxes, species ID, and estimated confidence value, and with a total count summary for the image at bottom left. Annotations include the primary species of interest; Salvin’s albatross (orange), and secondary species of interest; erect-crested penguins (green), and New Zealand fur seals (blue). Not shown in this image are fulmar prions (petrels)..

There was an exponential relationship between the amount of training data and the accuracy of the model represented by the F1 score (Fig 4), indicating that for an average multi-species model, at least 1000 examples of each target species would be required for successful training (leading to model success of >75%). In the current study, model success based on F1 scores were higher than this for albatross and penguins, reaching 92.9% and 89.4% success, respectively (Table 1, Fig 4).

**Fig 4.**
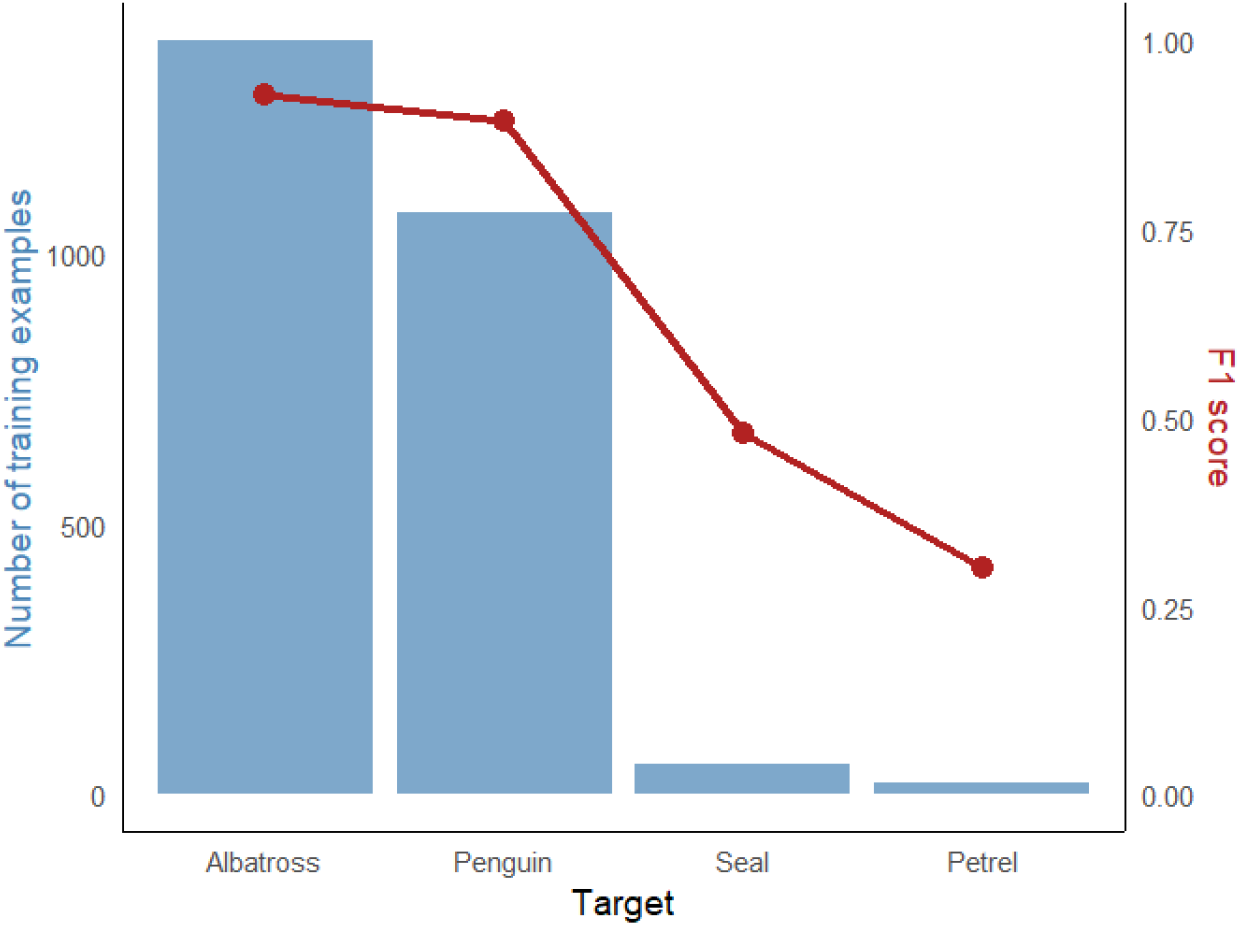
Plot of mixed-species model training prediction showing example species on the x-axis, and the number of training examples (blue bars, left axis) versus the F1 score (red line, right axis). A minimum of 1000 training examples were expected to be necessary to achieve an average model accuracy of >75% when adding a new species – This was achieved for albatross and penguins with initial training accuracy (F1 scores) of 0.928 and 0.894 respectively.

## Discussion

This study presents a novel technical method for considerably improving the efficiency of automated identification of animals from images, utilising a high accuracy AI solution. The model’s high initial detection accuracy (F1 score of 92.8%) for the target species (Salvin’s albatross) demonstrates the ability of the AI to recognise and count targets accurately. The results presented here are more accurate than similar studies, for example Rogers et al (2025) who reported average F1 scores ranging from 50.2 – 75.0% for AI counts of drone-based imagery on the same species at the same location. F1 scores can be influenced by the algorithm used by the model, the resolution of the images used for training and detection, and the assessment methodology, so these results therefore represent the accuracy of the end result. The F1 score also represents a minimum initial accuracy value of the model, which can be further improved by ongoing revision (by annotating any false positives or false negatives) each time the model is run. Revised training data can then be incorporated into the model to improve detection ability next time it is run. In addition to demonstrating high accuracy compared to other solutions, our AI model is simple and fast to train on new target types, adding value to conservation managers.

Based on the results from this study, training using a minimum of 1000 examples would be expected to result in model success of over 75% for a new species, (Fig. 4), which was threshold of accuracy considered acceptable for this project. The lower initial F1 scores observed for non-target species are a result of their lower numbers present in the training imagery (particularly in the case of seals and petrels) meaning the model did not receive sufficient training data to achieve optimal accuracy on those species. The detection ability for penguins was lower than for albatross, which is expected as the model was trained on fewer examples. Successful detection of penguins was also affected by the lower contrast between them and their surrounding substrate – they were commonly found in holes and crevices with deep shadows and often had their darker dorsal surface uppermost, making detection more difficult. Despite these difficulties, the relatively high initial accuracy (89.4%) achieved for this secondary species was in part due to the high resolution imagery utilised by the model, meaning that it was able to recognise the penguins’ yellow head crests to assist with identification – a feature which would not be possible with lower-resolution data. Additional training data would improve results still further, demonstrating the flexibility of the model to be quickly trained to recognise other target species.

In this study, the AI received some cursory training to recognise multiple secondary target species, allowing simultaneous counts not only of the species of interest (Salvin’s albatross), but also of other species at the Bounty Islands (erect-crested penguin, New Zealand fur seal, and petrel). All of the secondary species included less training data than the predicted minimum amount, so in all cases providing the model with additional training data would be expected to improve performance. Given that the primary focus of this research was on identifying and counting albatrosses, the ability to simultaneously survey multiple other species provides an added bonus for conservation management, and this can be improved on to provide simultaneous surveys of multiple species for the future. In addition to the species described here, the AI is already trained to recognise additional targets including various terrestrial species of birds and mammals, and also ships, with a view to future research identifying and counting other species. This would provide further use cases conducting breeding counts and population estimates for other species including breeding seabirds and pinnipeds, as well as investigating seabird-vessel interactions at sea.

In the original study based on manual counting, the raw count of Salvin’s albatross at the Bounty Islands was 60,419 birds, and after correction factors were applied to account for the proportion of birds loafing rather than occupying a nest, this was used to calculate the number of Apparently Occupied Nest Sites as 57,350. From this, a further correction factor was applied to account for the proportion of non-breeding birds, generating a population estimate of between 41,723 and 26,955 breeding pairs (Baker & Jensz, 2019). For the current study, not all of the original images were available for re-counting so the AI was presented with around half of the original image data (30,899 birds). From this, the AI re-count was 97% accurate with the original manual count (Table 3).

The purpose of the current study was to use an AI counting tool to produce a revised count. To derive a population estimate from these data it would be necessary to apply suitable correction factors to account for non-breeding birds, and those loafing or not on a nest when the image was taken. The AI re-count was slightly higher than the human count by 819 birds, or 3% (Table 3), which could be explained by the manner of counting. The human counter counted nesting birds only and ignored any flying birds as they could be the partner of a nesting bird already counted, or a non-breeding individual, and therefore would not provide a good indicator of the breeding population (Baker & Jensz, 2019). Conversely, this version pf the AI model was trained to count all the albatrosses in an image, which included flying birds, and would explain the higher overall count number. For the future, a revised version of the AI could be trained to differentiate between flying and nesting birds, with a view to generating a more accurate population estimate. In some images (e.g. Molly Cap and Funnel Is) the AI count was slightly lower than the human count. This is most likely due to mis-identification of an albatross as one of the secondary species, so this discrepancy could also be reduced with some additional revision and retraining of the model, particularly if there were specific factors in those images which affected detection accuracy, for example reduced contrast of the targets compared to the background, which could occur as a result of local topography or environmental conditions when the picture was taken.

Using a piloted aircraft for fast data collection in conjunction with an advanced AI counting model for efficient data processing allowed this population survey to be conducted in a single day. In comparison, ground counts of seabird breeding populations at similar isolated locations in the sub-Antarctic typically require field teams to be deployed on-site for weeks at a time (Muller et al., 2020), and the close approach of field crews can introduce disturbance to nesting birds and other non-target species. Imagery collected using UAVs can be restricted by the limited flight time of multi-rotor drones (Chrétien et al., 2016), so the versatile, high-resolution system presented here offers considerable advantages when searching larger, and more remote sites.

Drone surveys of sub-Antarctic seabird populations require a field team to travel to the island and typically remain there for at least a week while conducting multiple drone flights (Muller et al., 2019; Parker et al., 2017; Rogers et al., 2025), meaning the method we describe here represents a considerable saving in time and effort. With the assistance of our AI recognition tools we now have the ability to process data and count animals in just a few minutes, providing results immediately. These results demonstrate the ability to improve efficiency of future population censuses, and enhance conservation management outcomes.

